# A targeted genetic modifier screen in Drosophila uncovers vulnerabilities in a genetically complex model of colon cancer

**DOI:** 10.1101/2022.09.23.508936

**Authors:** Ishwaree Datta, Benjamin Linkous, Tyler Odum, Christian Drew, Andrew Taylor, Tajah Vassel, Erdem Bangi

## Abstract

Kinases are key regulators of cellular signal transduction pathways. Many diseases including cancer are associated with global alterations in protein phosphorylation networks, as a result, kinases are frequent targets of drug discovery efforts. However, target identification and assessment, a critical step in targeted drug discovery which involves identifying essential genetic mediators of disease phenotypes, can be challenging in complex, heterogeneous diseases like cancer where multiple concurrent genomic alterations are common. Drosophila is a particularly useful genetic model system to identify novel regulators of biological processes through unbiased genetic screens. Here, we report two classic genetic modifier screens focusing on the Drosophila kinome to identify kinase regulators in two different backgrounds: KRAS TP53 PTEN APC, a multigenic cancer model that targets four genes recurrently mutated in human colon tumors and KRAS alone, a simpler model that targets one of the most frequently altered pathways in cancer. These screens identified hits that are shared by both models as well as those unique to each one, emphasizing the importance of capturing the genetic complexity of human tumor genome landscapes in experimental models. Our follow-up analysis of two hits from the KRAS only screen suggest that classical genetic modifier screens in heterozygous mutant backgrounds that result in a modest, non-lethal reduction in candidate gene activity in the context of a whole animal —a key goal of systemic drug treatment— may be a particularly useful approach to identify most rate limiting genetic vulnerabilities in disease models as ideal candidate drug targets.

## INTRODUCTION

Cancer is a genetically complex and heterogeneous disease. For most solid tumors, tumorigenesis and progression into metastatic disease typically require the accumulation of multiple genomic alterations (Bailey *et al*. 2018; Sanchez-Vega *et al*. 2018). For instance, broad alterations in multiple signaling networks, global changes in transcriptomic, proteomic, phospho-proteomic and epigenomic profiles have consistently emerged from recent tumor profiling studies as hallmarks of most advanced tumors (Weber *et al*. 2020; Das *et al*. 2020; Zhang *et al*. 2022). Individual genes and pathways recurrently altered in human tumors have been well characterized in multiple experimental systems and multigenic cancer models have started to shed light into emergent interactions between concurrent cancer driving genetic alterations (Kersten *et al*. 2017; Guerin *et al*. 2020; Sajjad *et al*. 2021). However, the complex, multigenic nature of tumor genome landscapes makes it particularly challenging to identify genes that are required to establish and maintain tumor phenotypes and prioritize candidate targets for drug discovery approaches (Haley and Roudnicky 2020).

Drosophila, with its sophisticated genetic tools and practical advantages, has a strong track record as a disease model and has also emerged as a useful drug discovery platform (Cheng *et al*. 2018; Mohr and Perrimon 2019; Munnik *et al*. 2022). Disease models generated either by genetically manipulating Drosophila orthologs of human disease genes or directly introducing disease associated variants of human genes into Drosophila have provided important insights into molecular mechanisms underlying human diseases including cancer (Verheyen 2022). Furthermore, multiple studies over the past decade have demonstrated a high degree of conservation of compound activity in Drosophila (Su 2019; Munnik *et al*. 2022); several compounds identified through these studies have been effective in vertebrate experimental models and human patients (Dar *et al*. 2012; Bangi *et al*. 2016, 2019, 2021; Sonoshita *et al*. 2018; Das *et al*. 2018).

We have previously leveraged the genetic simplicity and power of Drosophila as a model system to functionally explore genome landscapes of human colon tumors (Bangi *et al*. 2016, 2019). These multigenic, tumor genome-based models capture many hallmarks of human tumors and demonstrate that intrinsic drug resistance is an emergent feature of genetic complexity. In this study, we take advantage of a classic genetic modifier screen approach to identify *bona fide* regulators of a 4-hit model targeting Drosophila orthologs of KRAS, TP53, PTEN and APC, four genes recurrently mutated in colon tumors, as well as KRAS alone, which represents one of the most commonly observed pathway alterations in human tumors (Cancer Genome Atlas Network 2012). As kinases are key regulators of signaling networks and frequent targets of cancer drug discovery efforts (Attwood *et al*. 2021), we focused our genetic screen to the Drosophila kinome, which consists of 377 kinases representing all branches of the human kinome tree (Manning *et al*. 2002; Gramates *et al*. 2022). Our parallel screens of these two models identified partially overlapping hits, indicating that as the genetic complexity of the model increases, its dependence on the activity of some kinases decreases and new genetic vulnerabilities that were absent in simpler models emerge. By comparing the sensitivity of our models to a genetic copy loss versus strong, tissue specific knockdown of kinase activity, we found that classic modifier screens in heterozygous mutant backgrounds, which result in a modest, non-lethal reduction of kinase activity in the whole animal may be a useful approach for identifying most rate limiting genetic vulnerabilities that could be exploited in therapy, while tissue specific gene knock-down or knock-out approaches can be used to uncover genetic dependencies.

## METHODS AND MATERIALS

### Drosophila strains

All Drosophila strains were maintained at room temperature on standard Drosophila medium. *w*^*1118*^ and all kinome stocks used for the screen were obtained from Bloomington Drosophila Stock Center (BDSC, see Supplemental Table 1 for details). TRIP RNAi lines (Zirin *et al*. 2020) UAS-akt^RNAi^ (BDSC #31701) and *UAS-dsor*^*RNAi*^ (BDSC #33639) used for *akt* and *dsor* knock-down were also obtained from BDSC. Transgenic Drosophila lines used to generate the KRAS TP53 PTEN APC combination are: *UAS-ras*^*G12V*^ (II, G. Halder) and three RNAi lines *UAS-p53*^*RNAi*^ (II), *UAS-pten*^*RNAi*^ (III) and *UAS-apc*^*RNAi*^ (II) that were obtained from the Vienna Drosophila Resource Center VDRC (Dietzl *et al*. 2007). The *UAS-ras*^*G12V*^*UAS-p53*^*RNAi*^ *UAS-pten*^*RNAi*^ *UAS-apc*^*RNAi*^ quadruple multigenic combination was generated by jumping the 3rd chromosome *UAS-pten*^*RNAi*^ transgene from VDRC onto the 2nd chromosome and recombining it into the previously described *UAS-ras*^*G12V*^*UAS-p53*^*RNAi*^ *UAS-apc*^*RNAi*^ triple combination on the second chromosome (Bangi *et al*. 2016). The *byn-gal4 UAS-GFP tub-gal80*^*ts*^ triple recombinant chromosome used for target expression in the hindgut has also been described previously (Bangi *et al*. 2012).

Fly lines that combine all transgenic elements required for targeted expression of KRAS TP53 PTEN APC or KRAS to generate the lethal screening phenotypes into a single genetic background (i.e. screening stocks) were generated using standard Drosophila genetic crosses: *w*^*1118*^ *UAS-dcr2/Y, hs-hid; UAS-ras*^*G12V*^ *UAS-p53*^*RNAi*^ *UAS-pten*^*RNAi*^ *UAS-apc*^*RNAi*^ ; *byn-gal4 UAS-GFP tub-gal80*^*ts*^ */S-T, Cy, tub-gal80, Hu, Tb* and *w*^*1118*^ *UAS-dcr2/Y, hs-hid* ; *UAS-ras*^*G12V*^ ; *byn-gal4 UAS-GFP tub-gal80*^*ts*^ */S-T, Cy, tub-gal80, Hu, Tb. UAS-GFP* and *UAS-dcr2* transgenes are included to fluorescently label targeted hindgut epithelial cells and to facilitate RNAi mediated knock-down, respectively. *tub-gal80* transgene (Lee and Luo 1999) introduced into the balancer chromosome inhibits gal4 activity in screening stocks to prevent transgene expression and lethality. *tub-gal80*^*ts*^ transgene on the 3rd chromosome, which drives the ubiquitous expression of a temperature sensitive allele of the Gal4 inhibitor Gal80 (McGuire *et al*. 2004), is used to temporally regulate transgene induction. The Y chromosome *hs-hid* transgene, which results in ubiquitous activation of apoptosis when induced (Starz-Gaiano *et al*. 2001), is used to kill all male progeny and facilitate mass virgin female collection required for the large number of crosses of the genetic modifier screens. Kinase alleles were balanced using balancer chromosomes carrying a Tb marker, using *CyO, Tb, RFP* (CTR) and *FM7c, Tb, RFP* (FTR) balancers for the second and X chromosomes respectively (Pina and Pignoni 2012) and the *TM6b, Hu, Tb* balancer for the third chromosome.

### Genetic screen

Mutant kinase alleles were introduced into the KRAS TP53 PTEN APC and KRAS backgrounds with a simple F1 cross (see Supplemental Figure 1 for examples). For the 2nd, 3rd and 4th chromosome screens, males from kinase stocks were crossed to virgin females from the KRAS TP53 PTEN APC and KRAS only screening stocks. For kinases on the X chromosome, *kinase*^*-*^*//FM7c, Tb, RFP* virgin females were crossed to males from the KRAS TP53 PTEN APC and KRAS only screening stocks. KRAS TP53 PTEN APC and KRAS virgins crossed to *w*^*1118*^/Y males (no kinase mutations) were used as our baseline controls for our screening read-outs. Virgins required for these crosses were generated *en masse* by two independent 1 hour heat shocks of screening lines using a 37^0^C water bath during development using the following schedule: 3-day egg lay at room temperature in bottles with standard Drosophila medium followed by removal of parents, first heat-shock of the progeny on day 4 and second heat shock on either day 5 or day 7.

**Figure 1.**
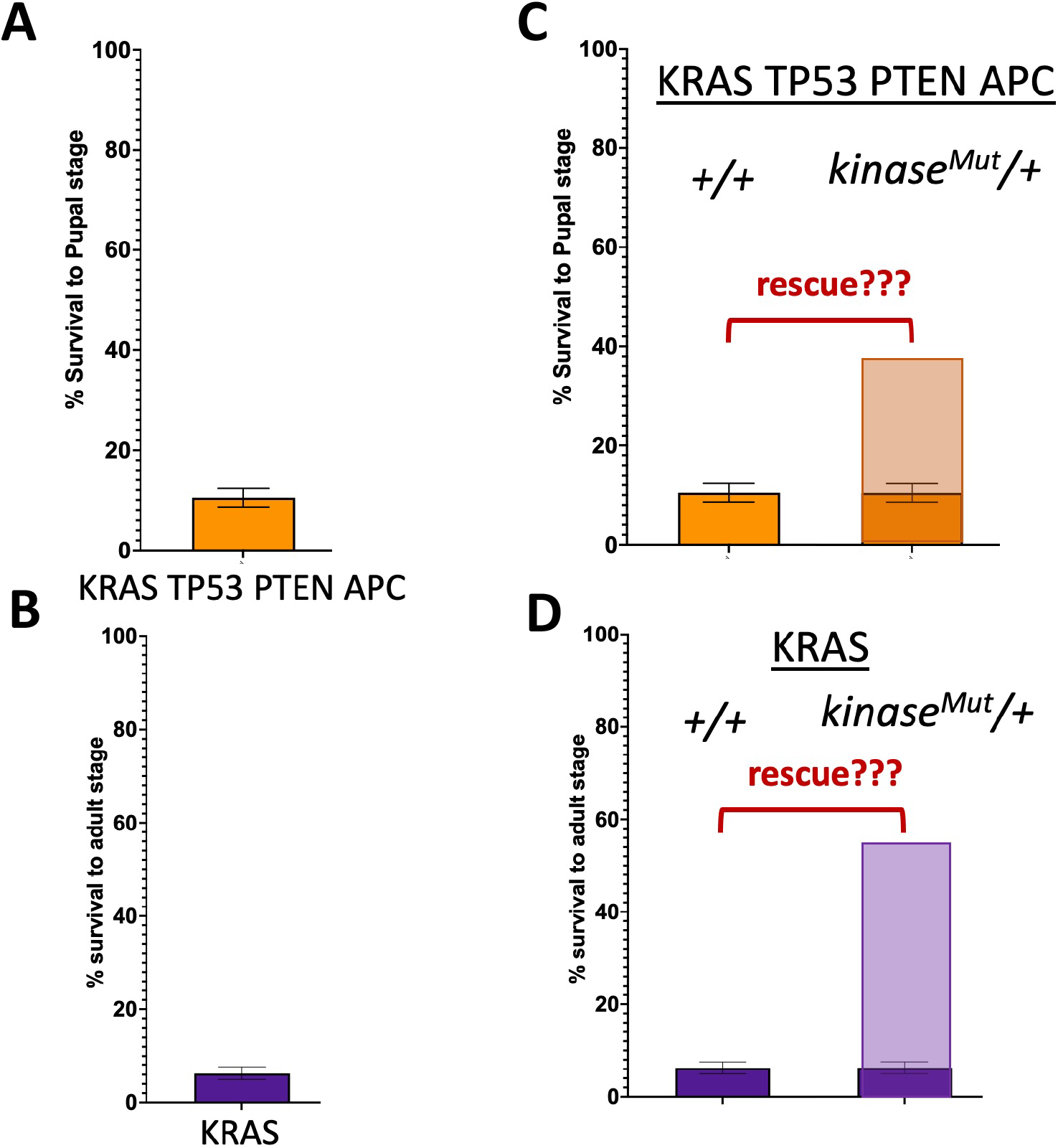
Screening strategy using rescue of KRAS TP53 PTEN APC and KRAS induced lethality as a primary read-out. (A,B) Lethal phenotypes induced by targeting *UAS-ras*^*G12V*^ *UAS-p53*^*RNAi*^ *UAS-pten*^*RNAi*^ *UAS-apc*^*RNAi*^ (KRAS TP53 PTEN APC, A) and *UAS-ras*^*G12V*^ (KRAS alone, B) to the developing hindgut using the hindgut specific *byn-gal4* along with a *UAS-GFP* transgene. (C,D) Screening strategy using rescue of lethality as a read-out. A mutant allele for each kinase is introduced into KRAS TP53 PTEN APC (C) and KRAS (D) backgrounds to evaluate their effect on lethality. Kinases with important, rate-limiting roles in these backgrounds may improve survival.

Crosses for the screen were set-up on Bloomington semi-defined medium (Bangi *et al*. 2019) at 29^0^C using 18-20 virgin females and 8-10 males for each cross. Parents were removed after 2 days of egg laying. Progeny were scored 12-14 days after crosses were set up. Experimental pupae were identified by the absence of the Tb marker. Survival to pupal stage was calculated by counting the number of experimental and control pupae in each vial (Figure S1). To calculate survival to adult stage, the fraction of empty experimental pupal cases (indicating an adult fly has emerged) was calculated. Each kinase allele was tested in triplicate. Kinase alleles that resulted in a statistically significant increase in survival compared to KRAS TP53 PTEN APC or KRAS alone (Figure 1C,D) were considered candidate hits (multiple t-tests with Holm-Sidak correction for multiple hypotheses, PRISM software). Candidate hits were then retested in an independent set of experiments and kinases that showed statistically significant rescue in the retest experiments were considered confirmed hits. Screening crosses that were inconclusive due to low n were retested similarly.

### Dissections, imaging and scoring

Crosses to analyze the overall size of the hindgut imaginal ring area were set up similarly on Bloomington semi-defined medium using 18-20 virgin females and 8-10 males for each cross, but at 18^0^C to prevent early larval lethality. Parents were removed after 3 days of egg laying and progeny were kept at 18^0^C for an additional 3 days before transgenes were induced by a temperature shift to 29^0^C. Experimental larvae were identified based on the absence of the Tb phenotype and hindguts from larvae at the late third instar stage were dissected 3 days after induction. Dissections were performed in phosphate buffered saline (PBS), hindguts were fixed in ice-cold 4% paraformaldehyde in PBS for 15 minutes at room temperature followed by 3 rinses and a 15-minute wash in PBS. Hindguts were mounted the next day on Vectashield with DAPI and imaged at 10X magnification, at 1.5 Zoom using Leica SPE DM6 confocal microscope using the 488 nm and 405 nm lasers for visualizing the GFP labeled hindgut epithelium and nuclei (DAPI), respectively. 10-12 hindguts of each genotype were imaged and the area of the imaginal ring region of each hindgut was quantified using Fiji Image J Software in pixels (Schindelin *et al*. 2012). KRAS TP53 PTEN APC and KRAS screening stocks crossed to *w*^*1118*^/Y, was used as positive controls and GFP-only expressing hindguts generated by crossing *w*^*1118*^ *UAS-dcr2* ; *byn-gal4 UAS-GFP tub-gal80*^*ts*^ */TM6b Hu, Tb* virgins to *w*^*1118*^/Y males served as negative controls. Experimental and control groups were compared using one-way ANOVA (PRISM software).

### Western blot analysis

Crosses to generate experimental larvae for dissections were set up as described in the previous section. Hindguts for protein extraction were dissected from late third instar larvae lacking the Tb marker after 3 days of induction. 10 hindguts for KRAS TP53 PTEN APC and 15 hindguts for KRAS alone along with matching GFP only control larvae were used for making protein lysates (3 biological replicates/genotype). Larval hindguts were homogenized with motorized pestle (at 10 second pulses) in ice cold lysis RIPA buffer (37.5μl, Sigma-Aldrich #R0278) fortified with protease and phosphatase inhibitors (Sigma-Aldrich #4693132001 and Millipore Sigma #524627). Lysates were then centrifuged at 4^0^C for 10 min at 13,000 rpm. 35μl of supernatants were transferred to a fresh tube, followed by addition of 12.5μl of Sample buffer (Bio-Rad #1610792) and 2.5μl of 20X Reducing Agent (Bio-Rad #1610792). The lysates were boiled for 10 minutes in a 100^0^C heat block, centrifuged at 4^0^C for 5 min at 13,000 rpm and 50 μl of supernatant were transferred to new tubes. Lysates were stored at − 80^0^C until use. For western blot analysis, proteins were separated using 4–12% Criterion™ XT Bis-Tris Protein Gels (Bio-Rad #3450125). Primary antibodies used were Rabbit anti-phospho-AKT (1:1000, Cell Signaling Technology #4060), Mouse anti-diphospho-Erk (1:1000 Sigma-Aldrich #M8159) and Mouse anti-Syntaxin as loading control (1:1000 Developmental Studies Hybridoma Bank #8C3). The secondary antibodies were Anti-rabbit IgG, HRP-linked Antibody (1:1000, Cell Signaling Technology #7074) and Anti-mouse IgG, HRP-linked Antibody (1:5000, Cell Signaling Technology #7076). Protein bands were developed with Immobilon Western Chemiluminescent HRP Substrate solutions (Millipore Sigma #WBKLS0050) and visualized under UV light using BioRad’s ChemiDoc MP™ Imaging Platform. Bands were quantified using Fiji Image J (Schindelin *et al*. 2012). Statistical analysis was performed with one way ANOVA using PRISM software.

## RESULTS AND DISCUSSION

### Screen Strategy

Rescue from lethality has been shown to serve as a useful read-out in high throughput chemical and genetic screens to identify novel regulators of disease relevant phenotypes (Rudrapatna *et al*. 2014; Levinson and Cagan 2016; Bangi *et al*. 2019). In order to generate a robust, lethal phenotype suitable for large scale screening, we used the *byn-Gal4* driver expressed in the hindgut epithelium (Takashima *et al*. 2008) to target our 4 hit model KRAS TP53 PTEN APC to the developing hindgut. This model comprises an oncogenic UAS-ras^G12V^ transgene and three UAS-RNAi transgenes to knock down Drosophila orthologs of tumor suppressors APC, TP53 and PTEN (Bangi *et al*. 2016). We found that targeting KRAS TP53 PTEN APC multigenic combination to the developing hindgut epithelium resulted in a highly penetrant and consistent early larval lethal phenotype (Figure 1A) while targeting KRAS only (UAS-ras^G12V^) to the same tissue resulted in a similarly robust lethal phenotype albeit at a later stage during late pupal development (Figure 1B). We then used these lethal phenotypes as surrogate read-outs in parallel genetic screens to evaluate vulnerabilities of genetically complex and simple models and identify genes with essential roles in intestinal transformation.

Kinases are important in governing cancer relevant pathways (Gross *et al*. 2015; Cicenas *et al*. 2018) therefore we focused our screens on the Drosophila kinome. Like in humans, most Drosophila loci are recessive, that is, they show no phenotype or lethality when a single genomic copy is removed. However, losing a copy of a gene that promotes oncogenic transformation and organismal lethality in the context of a multigenic model may prove rate limiting and improve survival. Therefore, we decided to adopt a classic genetic modifier screening approach by reducing the gene dosage of each kinase by introducing a mutant allele into our models using publicly available fly lines. Building on prior studies taking a similar approach (Levinson and Cagan 2016; Sonoshita *et al*. 2018), we reasoned that reducing the level of a kinase that may be critical for tumor progression would rescue the lethality we observe in our models (Figure 1C,D).

### Screen Design

The ability to perform complex genetic manipulations and carry out large scale in vivo screens are two key strengths of Drosophila as a model system. However, as our multigenic models are already genetically complex —for instance, experimental animals in our KRAS TP53 PTEN APC model carry 8 different transgenes— performing additional genetic manipulations in these backgrounds, especially in the context of a large-scale genetic screen, can be challenging. To address this problem, we established screening lines for KRAS TP53 PTEN APC and KRAS only by consolidating all transgenic elements required to generate each lethal screening read-out into a single genetic background (Figure S1). These screening lines also include 1) a ubiquitously expressed *gal80* transgene (Lee and Luo 1999) to prevent gal4 activity and lethality within screening stocks and 2) a *hs-hid* transgene on the Y chromosome (Starz-Gaiano *et al*. 2001) that leads to ubiquitous activation of apoptosis and lethality in male progeny when induced, a strategy that allows *en masse* virgin generation necessary for large scale screens. This design allowed us to introduce mutant kinase alleles into KRAS TP53 PTEN APC and KRAS alone backgrounds with simple F1 genetic crosses (Figure S1).

### Screen Results

The Drosophila kinome comprises 377 kinases (Gramates *et al*. 2022). We were able to obtain mutant lines for 206 kinases, covering 54.6% of the entire fly kinome (Table S1). Our parallel screens of the same set of kinases identified 3 confirmed hits from the KRAS TP53 PTEN APC screen (*aPKC, par-1, cdk2*), while KRAS screen resulted in 5 hits (*par-1, cdk2, hppy, dsor1, akt*) (Figure 2A-C).

**Figure 2:**
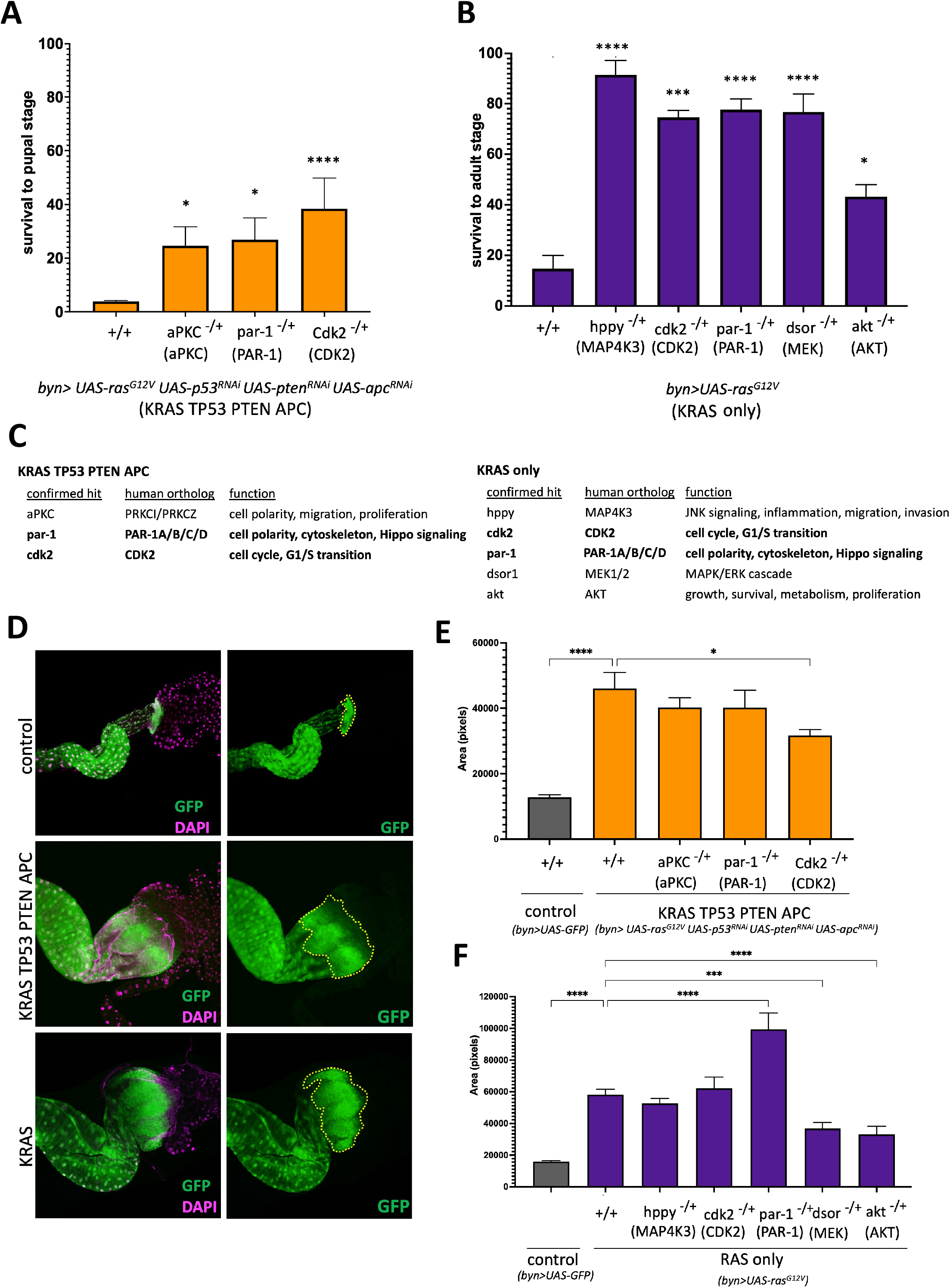
Hit confirmation and subsequent analysis. (A,B) Rescue of KRAS TP53 PTEN APC induced larval lethality (A) and KRAS induced pupal lethality (B) by confirmed hits from the kinome screens. Human orthologs are in parentheses. Error bars represent Standard Error of the Mean (SEM). * p≤0.05, *** p≤0.001, **** p≤0.0001 (multiple unpaired t-tests with Holm-Sidak Correction, PRISM Software). (C) Confirmed hits from KRAS TP53 PTEN APC (left) and KRAS (right) screens, with their human orthologs and biological functions. (D) Anterior regions of dissected control (GFP only), KRAS TP53 PTEN APC and KRAS alone hindguts. Hindgut epithelium is in green and nuclei in cyan. (E, F) Quantification of the imaginal ring area (outlined in yellow dashed lines in panels on the right in D) in hindguts with indicated genotypes, measured in pixels. Error bars represent Standard Error of the Mean (SEM). * p≤0.05, *** p≤0.001, **** p≤0.0001 (one way ANOVA, PRISM Software). Mutant alleles: *aPKC*^*k06403*^, *par-1*^*k06323*^, *cdk2*^*2*^, *hppy*^*SH1261*^, *dsor*^*LH110*^, *akt*^*04226*^

The two shared hits between the two screens were *par-1* and *cdk2*, both with highly conserved human orthologs and cancer relevant functions (Figure 2C). *cdk2* regulates G1/S transition of cell cycle in association with *CycE* and plays an important role in modulating cell division and differentiation (Sauer *et al*. 1995; Roesley *et al*. 2018). The human cdk2 ortholog, CDK2, is altered in various cancers including colorectal, and its downregulation can have anti-tumor effects depending on the genetic context (Tadesse *et al*. 2020). *par1*, the second shared hit, is a critical regulator of cytoskeleton structuring, cell polarity and Hippo signaling (Huang *et al*. 2013; Doerflinger *et al*. 2022). *par1* is the single Drosophila ortholog of PAR-1/MARK, a serine/threonine kinase family in mammals (Wu and Griffin 2017). PAR-1 has been implicated in cancer progression and is upregulated in various tumors like pancreatic, breast and lung cancer (Boire *et al*. 2005; Cisowski *et al*. 2011; Tekin *et al*. 2018). Hits like these are significant as potential drug targets as they may represent broad genetic sensitivities shared among different cancer genome landscapes.

*aPKC*, the only hit that was unique to KRAS TP53 PTEN APC (Figure 2A,C), is an atypical protein kinase with a large number of downstream targets and a key regulator of cell polarity (Vorhagen and Niessen 2014; Hong 2018). *aPKC* overexpression is also linked to progression and development of different cancers (Eder *et al*. 2005; Regala *et al*. 2005; Nayak *et al*. 2019). Of note, even though both *aPKC* and *par1* play important roles in cell polarity, only *par1* was a hit in both screens. It is possible that even though the same biological process is important in both genetic contexts, sensitivity of different models to a reduction in the activity of genes involved in that process may be different. It is also possible that the role of aPKC in the KRAS TP53 PTEN APC is unrelated to its role in cell polarity. Either way, the ability of this approach to identify context dependent requirements for the same gene in different genetic backgrounds illustrates its utility in identifying new leads and testable mechanistic hypotheses.

The three genes identified as unique hits from the KRAS only screen were *hppy, akt* and *dsor* (Figure 2B,C), which are key components of Hippo, PI3K and MAPK signaling cascades, respectively (Zheng *et al*. 2015; Yang *et al*. 2019; Guo *et al*. 2020). Hppy, a serine threonine kinase, activates Warts alongside Hippo and regulates JNK signaling, inflammation, cellular migration and invasion (Zheng *et al*. 2015; Meng *et al*. 2015; Liu *et al*. 2017; Chuang *et al*. 2018, 2019). Its human ortholog, MAP4K3, promotes cell migration and invasion in various cancers (Varghese *et al*. 2016; Hsu *et al*. 2016; Ho *et al*. 2016; Liu *et al*. 2017). *dsor*, the Drosophila ortholog of MEK1/2, is a central component of the MAPK signaling pathway which phosphorylates ERK and regulates several cancer relevant processes including cell growth, proliferation, differentiation, migration, apoptosis. Aberrant activation of MAPK signaling in tumors results in unregulated cell proliferation and downregulation of antiproliferative genes (Chambard *et al*. 2007; Guo *et al*. 2020). The third hit, *akt*, lies downstream of PI3K/AKT signaling cascade, another pathway that regulates cell growth, cell survival, proliferation and metabolism both in Drosophila and mammals (Saxton and Sabatini 2017) and is dysregulated in various cancers (Yang *et al*. 2019). Our observation that KRAS TP53 PTEN APC is not sensitive to a reduction the gene dosage of these kinases is interesting, particularly in the case of *dsor* and *akt*, given the strong activation of both MAPK and AKT signaling pathways in RAS-PI3K co-activated tumors (Wee *et al*. 2009). Overall, our findings suggest that as the genetic complexity of the model increases, it may become less dependent on kinases that have important roles in simpler genetic backgrounds.

### Characterizing hits using cancer relevant secondary assays

While rescue from lethality is a useful surrogate screening read-out (Rudrapatna *et al*. 2014; Levinson and Cagan 2016; Bangi *et al*. 2019), it is important to establish more disease relevant secondary assays to test whether hits from such screens can also modify disease phenotypes and explore their mechanisms of action. Use of multiple phenotypic assays for subsequent studies is particularly important for a complex disease like cancer. We have previously demonstrated that multigenic cancer models in Drosophila capture multiple key hallmarks of human tumors (Bangi *et al*. 2012, 2019, 2021) and here we use one such assay to test our hits from the genetic screens as a proof of concept.

Targeting KRAS TP53 PTEN APC and KRAS to the larval hindgut epithelium results in the expansion of hindgut imaginal ring zone (Figure 2D), the anterior region of the hindgut where proliferating progenitor cells reside (Murakami and Shiotsuki 2001). We next tested whether hits from our screens could reduce KRAS TP53 PTEN APC or KRAS-induced expansion of this region as a secondary read-out. We found that a reduction in *cdk2* resulted in a significant decrease in the size of the imaginal ring area in KRAS TP53 PTEN APC background but not in KRAS alone (Figure 2E,F). We also found that reduction in *akt* or *dsor* levels resulted in a significant reduction in the imaginal ring zone size in KRAS alone (Figure 2F), further confirming the important roles these two kinases play as downstream mediators of the RAS pathway. Finally, *par1* heterozygosity resulted in a notable increase in the size of the larval hindgut imaginal ring region in KRAS (Figure 2F). As the size of this area is likely to be determined by a combination of proliferation rates, apoptosis, cell size and cell polarity, subsequent assays that quantitatively analyze these markers will be necessary to investigate the mechanistic bases for these observations.

Combined, these results show that not all hits that rescued KRAS TP53 PTEN APC or KRAS induced lethality were able to reduce the size of the transformed hindgut area, suggesting that the expansion of the imaginal ring area cannot fully account for the lethal phenotype we observe in our models. These results emphasize the importance of using multiple assays that capture different hallmarks of tumors to fully explore mechanisms of action of hits generated from genetic screens.

### Tissue specific knockdowns uncover genetic dependencies on *dsor* and *akt*

Ras/MAPK and PI3K/AKT signaling cascades are two of the most common pathways dysregulated in human cancers including colorectal cancer and multiple experimental models including our previous work (Ericson *et al*. 2010; Bangi *et al*. 2016, 2019; Reggiani Bonetti *et al*. 2018; Jiang *et al*. 2020; Chen *et al*. 2021), making them important drug targets in cancer therapy. Still, we found that reducing *dsor* or *akt* gene dosage using loss of function alleles is not sufficient to rescue KRAS TP53 PTEN APC induced lethality in our screen, suggesting that this model was not very sensitive to changes in *akt* or *dsor* levels. To investigate the genetic dependency of our multigenic model on these two genes, we tested the effect of strongly knocking down each gene in the hindgut on KRAS TP53 PTEN APC induced lethality. RNAi mediated knockdown of *dsor* or *akt* in the hindgut epithelium significantly improved survival to pupal stage in KRAS TP53 PTEN APC background (Figure 3A,C) and reduced the expansion of the imaginal ring area (Figure 3E,G). Consistent with results of our KRAS only screen, RNAi knock-down of *akt* and *dsor* also rescued KRAS induced lethality and expansion of the imaginal ring area (Figure 3B,D,F,H). These results demonstrate that even though both *akt* and *dsor* are required for intestinal transformation in KRAS TP53 PTEN APC background, this model is more resistant to moderate changes in *dsor* or *akt* levels and a stronger reduction in their activity is necessary to lead to a rescue. This is in contrast to the KRAS only background, where reducing the *akt* or *dsor* gene dosage is sufficient to mount a rescue.

**Figure 3.**
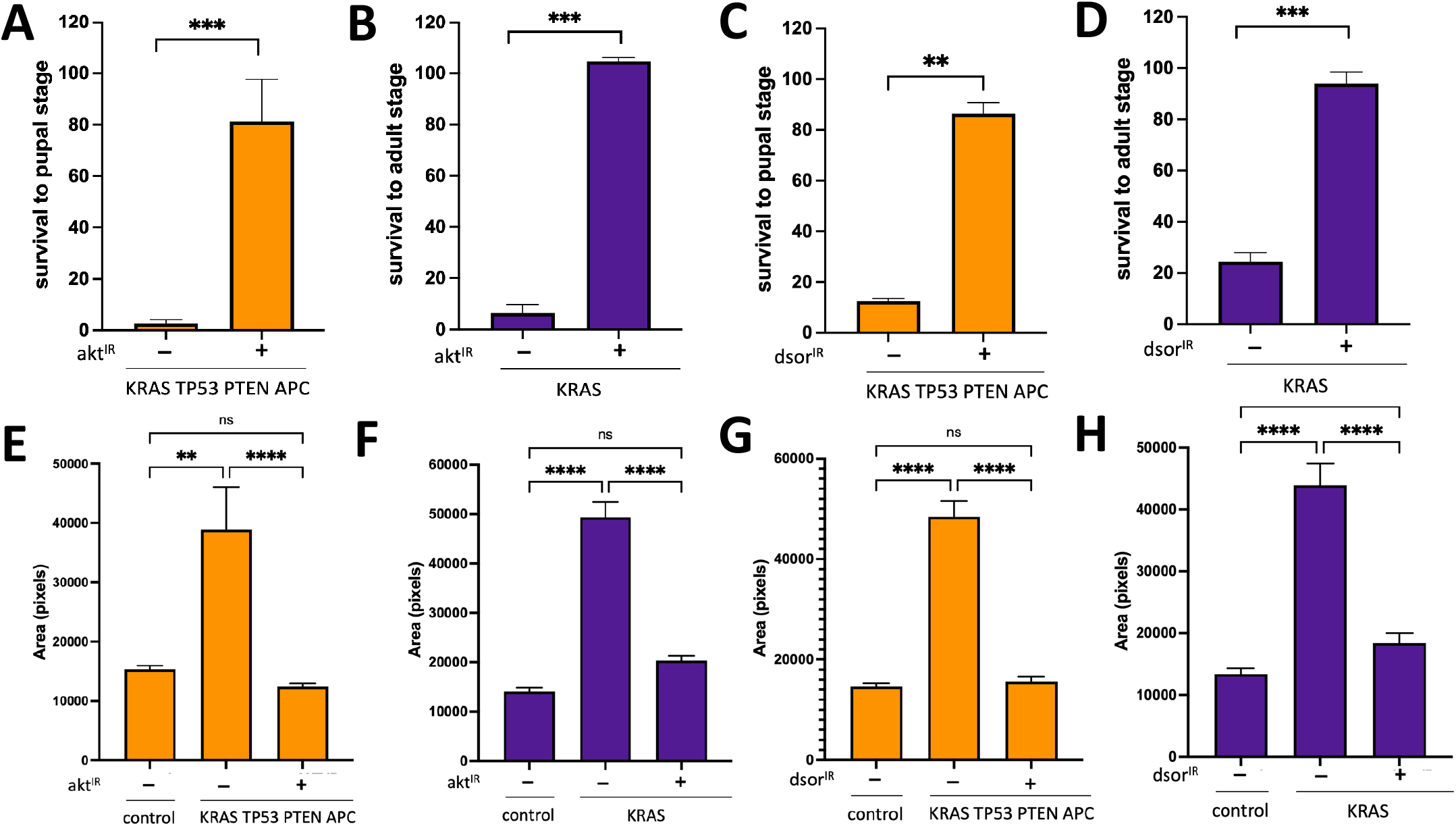
Strong-tissue specific knock-downs reveal requirements for both *dsor* and *akt* in KRAS TP53 PTEN APC. (A-D) Rescue of KRAS TP53 PTEN APC induced larval lethality (A,C) and KRAS induced pupal lethality (B,D) by knocking down *akt* (A,B) or *dsor* (C,D) in the hindgut epithelium. Error bars represent Standard Error of the Mean (SEM). ** p≤0.01, *** p≤0.001 (t-tests, PRISM software). (E-H) Quantification of hindgut imaginal ring area in hindguts with indicated genotypes. Error bars represent Standard Error of the Mean (SEM). ** p≤0.01, **** p≤0.0001 (one way ANOVA, PRISM Software).

Identifying genetic dependencies of tumors is an important tool for target identification and prioritization in drug discovery approaches. Our findings demonstrate that genetically complex models may not be sensitive to moderate reductions in the activity of all genes they are dependent on. This has important implications for target prioritization for drug discovery as it may not always be feasible to achieve a strong enough reduction in target activity comparable to what we observe in our tissue specific knockdowns by systemic drug treatment without significant side effects. As genetic modifier screens performed in heterozygous genetic backgrounds select for hits that suppress disease phenotypes without causing any organismal lethality by design, they may represent a useful approach to identify most rate limiting genetic vulnerabilities that may be more suitable candidate drug targets.

### Molecular correlates of genetic dependency and sensitivity to *akt* and *dsor*

To further investigate how reducing the gene dosage of *akt* or *dsor* and their RNAi knockdowns alter Ras/MAPK and AKT pathway activity in our models, we directly analyzed the levels of phoshpo-Akt (pAKT) and diphospho-Erk (dpERK), the main read-outs of the Ras/MAPK and AKT pathways, respectively in KRAS TP53 PTEN APC and KRAS only hindguts (Figure 4). Heterozygous loss of *akt* or *dsor* resulted in modest but statistically significant downregulation of dpERK in KRAS TP53 PTEN APC hindguts but had no significant effect on pAKT levels (Figure 4A,C,E), suggesting that increased AKT activity may be more critical than ERK activity in the context of KRAS TP53 PTEN APC. *dsor* knockdown in KRAS TP53 PTEN APC hindguts was not sufficient to alter pAKT levels but resulted in a much stronger reduction in pERK levels, indicating that KRAS TP53 PTEN APC is still dependent on ERK activity and can be rescued if a strong enough reduction in its activity is achieved (Figure 3C,G). *akt* knock-down on the other hand resulted in a significant reduction of both pAKT and pERK levels, albeit much stronger reduction in pAKT compared to dpERK (Figure 4C,E). These results demonstrate that in KRAS TP53 PTEN APC, a strong reduction in either AKT or MAPK pathway activity is sufficient to result in rescue, however, this level of reduction in activity could not be achieved in heterozygous mutant backgrounds for either *akt* or *dsor*.

**Figure 4.**
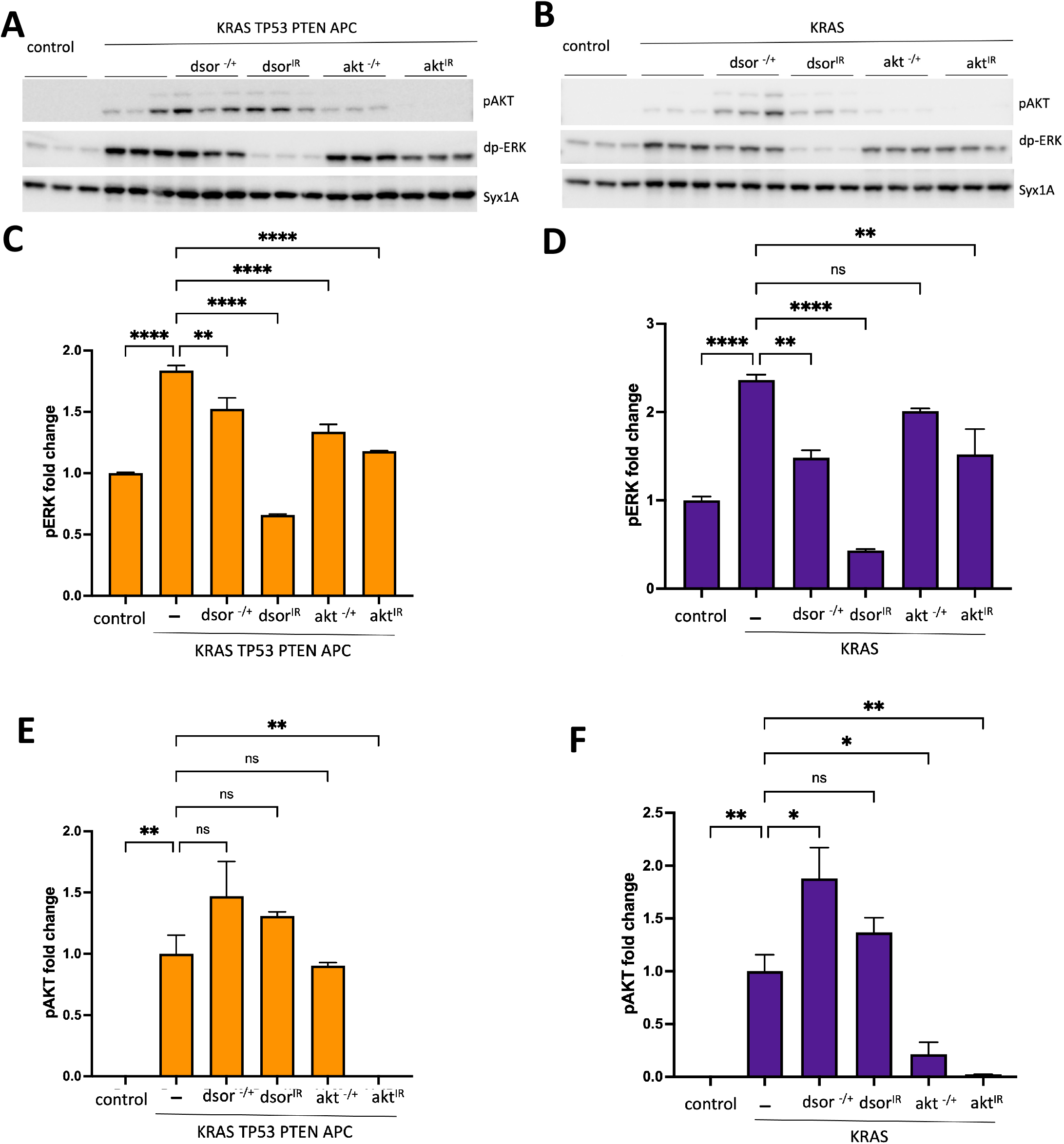
Analysis of MAPK and AKT pathway activity in response to tissue specific knock-down versus heterozygous loss. (A,B) Western blot analysis of pAKT and pERK levels in protein extracts from hindguts with indicated genotypes (3 biological replicates/genotype). Syntaxin: loading control. (C-F) Quantification of western blot data presented in (A,B). For quantification of pERK levels, all genotypes are normalized to GFP-only negative controls (C,D). As there were no detectable pAKT bands in GFP-only negative controls (top blots in A,B), all genotypes were normalized to KRAS TP53 PTEN APC (E) or KRAS (F). Error bars represent Standard Error of the Mean (SEM). *p≤0.05, ** p≤0.01, **** p≤0.0001 (one way ANOVA, PRISM Software).

In comparison, in KRAS only hindguts, we observed a more significant decrease in MAPK and AKT pathway activities in *dsor* and *akt* heterozygous mutant backgrounds, respectively (Figure 4B,D,F). We also noted a significant increase in pAKT levels in *dsor* heterozygous mutant background, most likely due to the disruption of a previously established negative feedback regulation of AKT activity by MEK (Turke *et al*. 2012). Our finding that heterozygous *dsor* loss is sufficient to rescue both KRAS induced lethality and imaginal ring area expansion despite an increase in pAKT levels suggests that in a KRAS only background, transformed tissue is more dependent on MAPK activity than AKT signaling; whereas KRAS TP53 PTEN APC transformed tissue is more dependent on AKT pathway. Combined, these results demonstrate that strong downregulation of either MAPK and AKT signaling pathways results in rescue of lethality and significant decrease in size of the transformed tissue (Figure 3), demonstrating that both pathways are essential for KRAS TP53 PTEN APC and KRAS alone. However, it is easier to achieve a level of reduction in MAPK or AKT pathway activity sufficient to rescue lethality or overgrowth in KRAS only compared to KRAS TP53 PTEN APC.

## Concluding remarks

AKT and MEK have been particularly attractive targets for cancer drug discovery given their central roles in Ras/MAPK and PI3K/AKT pathways and functional studies demonstrating a requirement for these two genes in tumor progression and development (Yap *et al*. 2011; Caunt *et al*. 2015; Saura *et al*. 2017; Bose and Kalinsky 2021). Despite extensive efforts, single agent MEK and AKT inhibitors have not been very successful in clinical trials for most solid tumor types in part due to severe toxicities (Chandarlapaty *et al*. 2011; Serra *et al*. 2011; Yap *et al*. 2011; Zhan *et al*. 2019; Faião-Flores *et al*. 2019). In addition, moderate inhibition of the MAPK pathway by MEK inhibitors can also lead to the development of resistance mainly via upregulation of PI3K/AKT pathway (Balmanno *et al*. 2009; Turke *et al*. 2012; Irvine *et al*. 2018; Tsubaki *et al*. 2019). Importantly, AKT and MEK inhibitor combinations designed to inhibit both pathways simultaneously and prevent the activation of feedback loops that promote resistance have also been unsuccessful due to adverse effects that were more severe than single agent trials and observed at lower doses (Tolcher *et al*. 2015, 2020). Overall, our findings are consistent with clinical studies suggesting that despite the importance of these genes in cancer, pharmacological inhibition of their activity to a level that could be detrimental to genetically complex tumors may not be possible in most cases due to significant side effects and toxicities (Altomare *et al*. 2010; Ma *et al*. 2017; Greenwell *et al*. 2017; Mondaca *et al*. 2018; Glen *et al*. 2022).

As most drugs are administered systemically, identifying doses that provide clinical benefit without any adverse effects and toxicities can be challenging. (Muller and Milton 2012). In fact, a large fraction of drugs that enter clinical trials fail due to unmanageable toxicities or lack of clinical efficacy at maximum tolerable doses even though they target key cancer genetic dependencies (Sun *et al*. 2022). While tissue specific gene knock-down and knock-out approaches are useful to identify genes essential for mediating disease phenotypes, genetic dependency of a disease on a gene does not necessarily reflect its potential as a drug target. Our results suggest that classic genetic modifier screens performed in heterozygous mutant backgrounds that result in a modest, non-lethal reduction in gene activity in the context of a whole animal may be a particularly useful complementary strategy to identify promising genetic vulnerabilities that can be successfully exploited by therapy.

## FIGURE LEGENDS

**Supplementary Figure 1.**
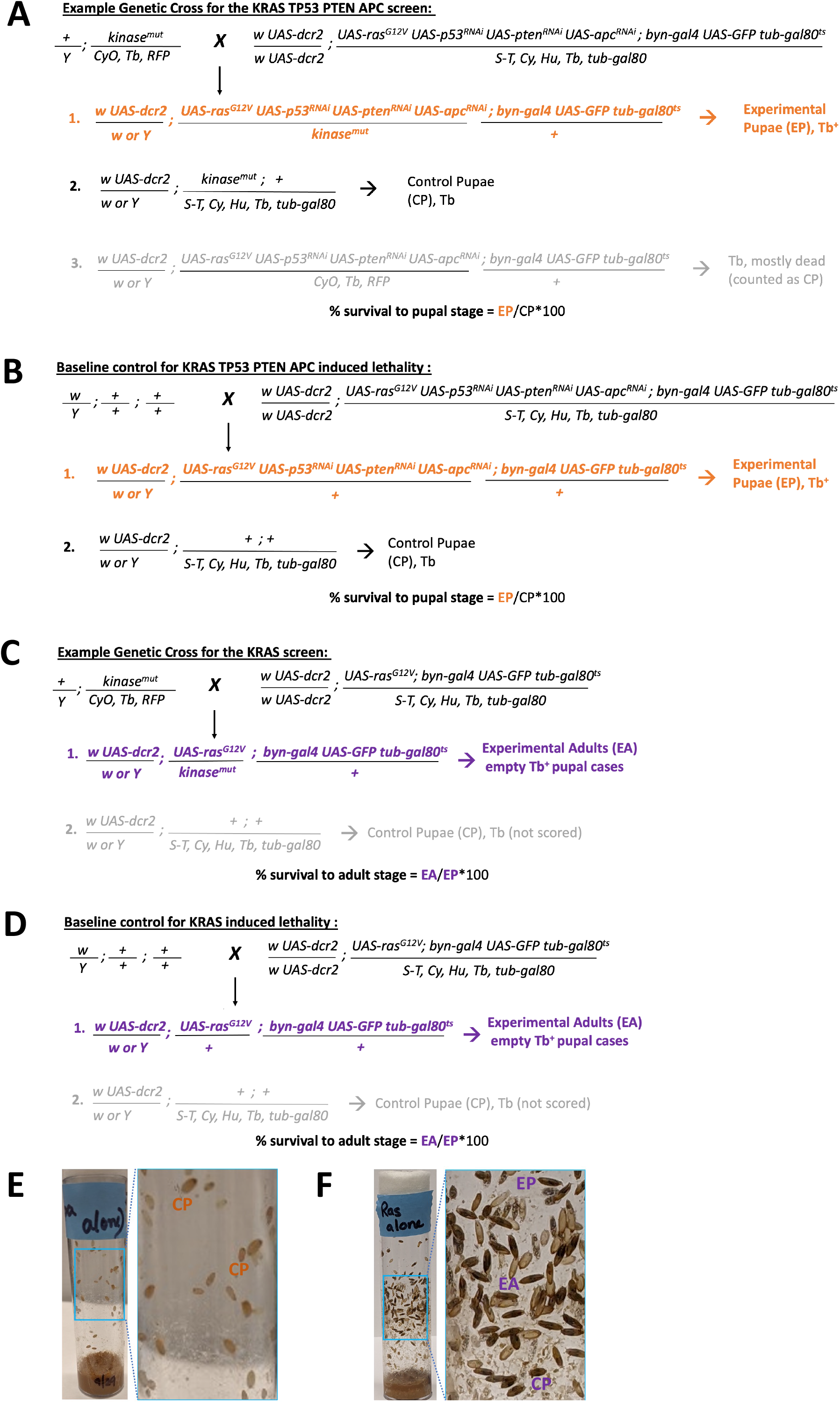
Genetic crosses for the screens. (A-D) Example crosses for KRAS TP53 PTEN APC (A,B) and KRAS (C,D) screens for kinases on the second chromosome (see methods for additional details). Experimental Pupae (EP) and Control Pupae (CP) were identified by the absence of the Tb marker. Experimental Adults (EA) were scored by identifying empty experimental pupal cases, indicating an adult fly has emerged. (E,F). Images of vials from the screens that show example EP’s, CP’s and EA’s.

## DATA AVAILABILITY STATEMENT

Drosophila fly stocks generated through this work are available upon request. The Bloomington Drosophila Stock Center (BDCS) ID’s and full genotypes of the kinome stocks used in this study are available in Supplementary Table 1 and online at https://bdsc.indiana.edu/index.html. More information on the kinase alleles used in this study can be found online at https://flybase.org/

## ACKNOWLEDGEMENTS

Stocks and transgenic TRIP RNAi lines (Office of the Director R24 OD030002: “TRiP resources for modeling human disease”) obtained from the Bloomington Drosophila Stock Center (NIH P40OD018537) and transgenic fly stocks obtained from the Vienna Drosophila Resource Center (VDRC, www.vdrc.at) were used in this study. We also thank Jason Cassara, Juan Martin Portilla, Justin Brown and Dr. Fateme Karimi Dermani for technical assistance.

---

**Supplemetary Table 1.**
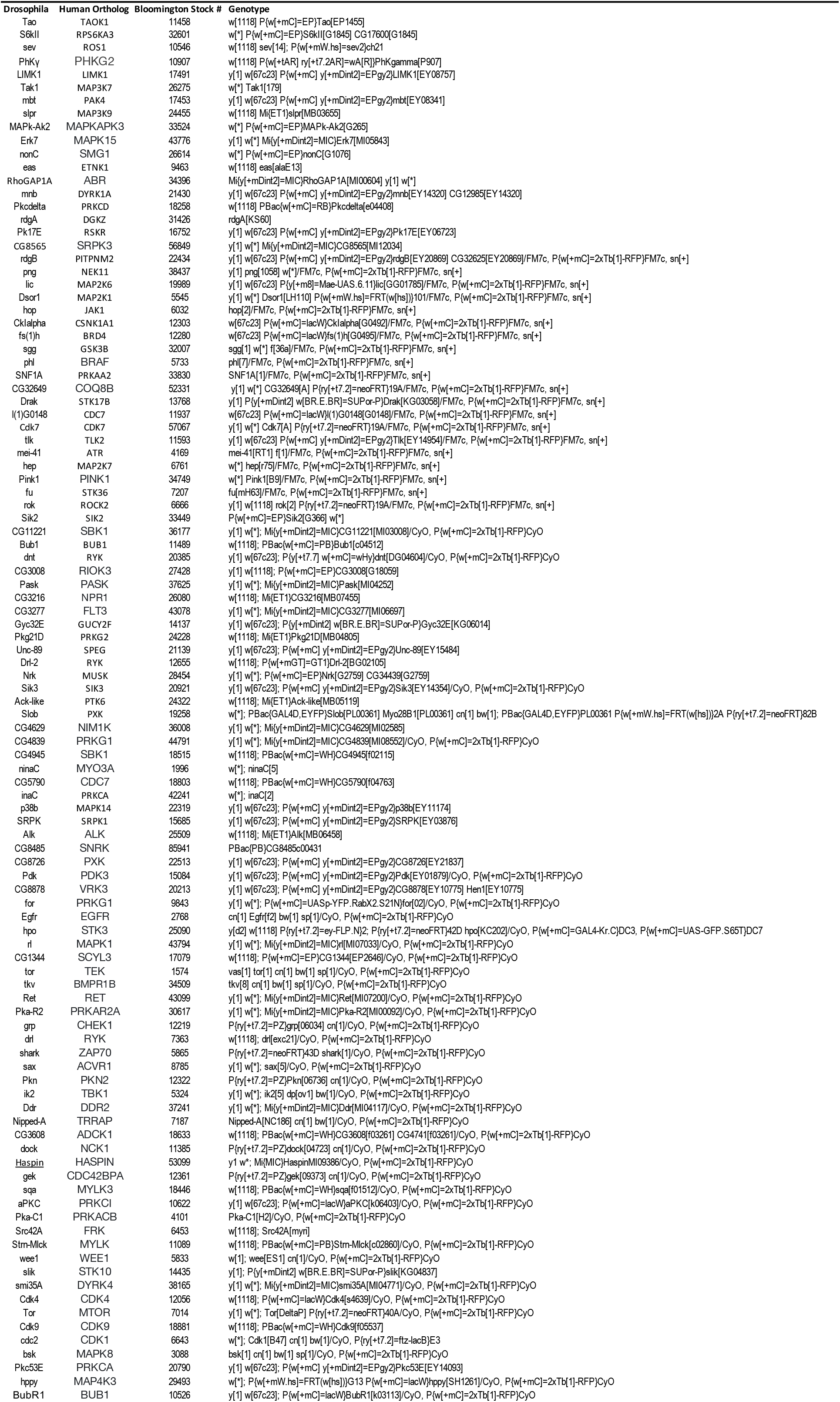
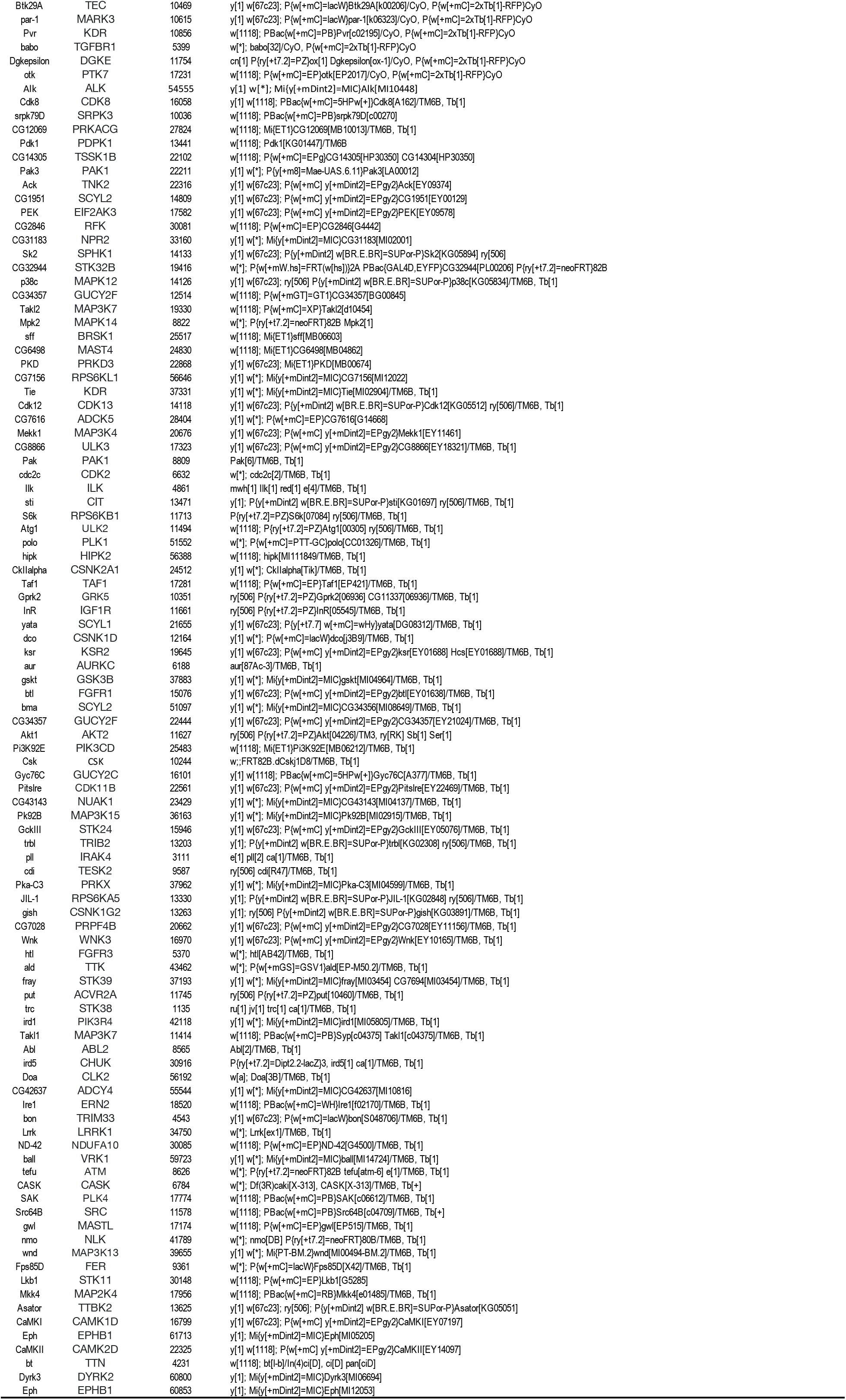
The kinome collection used in the screens.

## Notes

### Competing Interest Statement

The authors have declared no competing interest.

